# Binding of SARS-CoV-2 non-structured protein 1 to 40S ribosome inhibits mRNA translation

**DOI:** 10.1101/2023.02.24.529933

**Authors:** Hung Nguyen, Hoang Linh Nguyen, Mai Suan Li

## Abstract

Experiments have shown that non-structural protein 1 (NSP1) of SARS-CoV-2 is a factor that restricts cellular gene expression and prevents mRNA translation in the ribosome 40S subunit. However, the molecular mechanism of this phenomenon remains unclear. To clarify this issue, all-atom steered molecular dynamics and coarse-grained alchemical simulations were used to compare the binding affinity of mRNA to 40S ribosome in the absence and presence of NSP1. We found that NSP1 binding to the 40S ribosome dramatically increases the binding affinity of mRNA, which, in agreement with experiment, suggests that NSP1 can stall mRNA translation. The mRNA translation has been found to be driven by electrostatic mRNA-40S ribosome interactions. Water molecules have been demonstrated to play an important role in stabilizing the mRNA-40S ribosome complex. The NSP1 residues that are critical in triggering a translation arrest have been identified.

---

Severe acute respiratory syndrome coronavirus 2 (SARS-CoV-2) caused the 2019 coronavirus disease (Covid-19) worldwide pandemic, which affected millions of people.^1^ Like other coronaviruses, SARS-CoV-2 is an enveloped, positive-sense, single-stranded RNA virus, and its closely related phylogenetic species are known to infect a large number of vertebrate species.^2,3^ The SARS-CoV-2 genome consists of about 30 kb linear, one of the 5’-capped and 3’-polyadenylated RNA genomic components that make up coronavirus particles, encoding two large overlapping open reading frames in gene 1 (ORF1a and ORF1b), and includes various structural and non-structural proteins at the 3’ end.^4,5^ After entering host cells, the viral genomic RNA is translated by the cellular protein synthesis machinery to produce a set of non-structural proteins that render cellular conditions favorable for viral infection and viral mRNA synthesis.^6,7^ In cells infected with SARS-CoV-2, one of the most enigmatic viral proteins is a host shutoff factor called non-structural protein 1 (NSP1).^8^ NSP1 is the product of the N-terminus of the first open reading frame ORF1a and serves to suppress host gene expression and host immune response. Generally, NSP1 plays an important role in the viral life cycle.^9^

All viruses rely on cellular ribosomes for their protein synthesis and compete with endogenous mRNA for access to a translation machinery known as protein synthesis, which acts as a focal point of control.^10^ Host gene expression is limited by the common viral strategy of shifting translational resources toward viral mRNA.^11,12^ This phenotype termed host shutoff, increases the access of viral transcripts to ribosomes and promotes innate immune evasion.^11^Host shutoff is a hallmark of coronavirus infection and has significantly contributed to the suppression of innate immune responses in multiple pathogenic coronaviruses, including SARS-CoV identified in China in 2013, Middle East respiratory syndrome coronavirus, and pandemic SARS-CoV-2.^13–15^ SARS-CoV-2 induced host shutoff, which is multifaceted and involves inhibition of host mRNA splicing by NSP16, restriction of cellular cytoplasmic mRNA accumulation and translation by NSP1, and disruption of protein secretion by NSP8 and NSP9.^16–19^

NSP1 of SARS-CoV (SARS-CoV NSP1) and of SARS-CoV-2 (SARS-CoV-2 NSP1) bind to the 40S ribosomal subunit and stall canonical mRNA translation at various stages during initiation.^20,21^ However, although *in vitro* binding and translation assays revealed that both SARS-CoV NSP1 and SARS-CoV-2 NSP1 exert similar efficacy in the host translational shutdown mechanism,^22^ SARS-CoV-2 was shown to be more infectious and triggers more co-morbid conditions than SARS-CoV.^23,24^ Binding of NSP1 to the ribosome results in endonucleolytic cleavage and subsequent degradation of the host mRNA. Importantly, interactions between NSP1 and a conserved region in the 5’ untranslated region of viral mRNA prevent the shutdown of viral protein expression through an unclear mechanism (Figure 1A). Thus, NSP1 inhibits all mechanisms of cellular antiviral defense that depend on the expression of host factors, including the interferon response. This shutdown of key parts of the innate immune system may facilitate efficient viral replication^25,26^ and immune evasion. Its key role in attenuating the antiviral immune response makes NSP1 a potential therapeutic target.^27,28^

**Figure 1:**
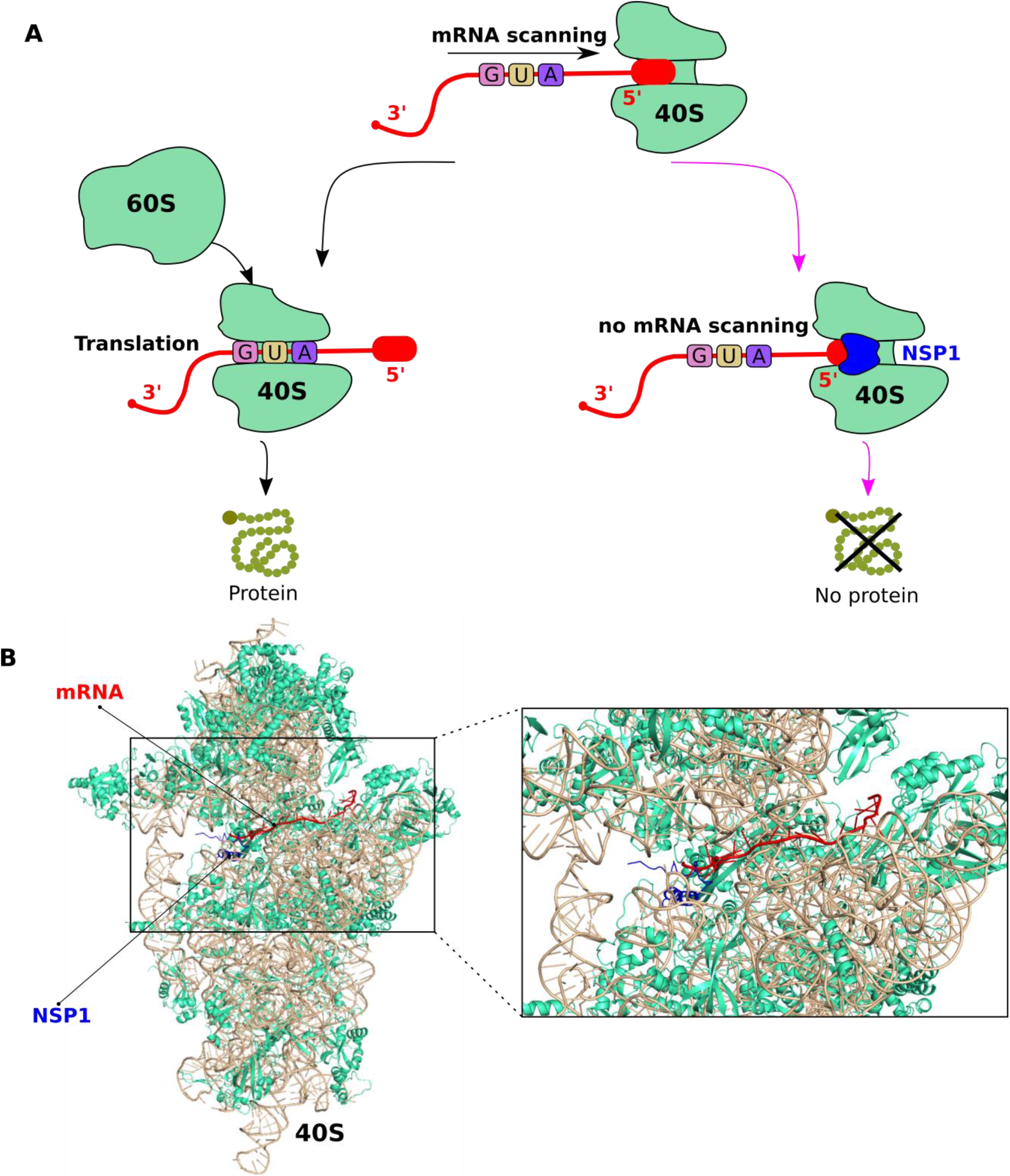
(A) Scheme of NSP1 action to suppress mRNA translation. (B) 3D structure of the mRNA- 40S-NSP1 complex constructed from a superposition of two different PDB structures 6ZOJ and 6HCJ. This structure includes ribosomal RNA (wheat), NSP1 (blue), mRNA (red), and ribosomal proteins (green-cyan).

Recently, Borišek *et al*.^29^ used all-atom simulation to investigate the interaction between NSP1 and the 40S subunit of the ribosome. They found that upon NSP1 binding to 40S, the critical switch of Gln158/Glu158 and Glu159/Gln159 residues of SARS-CoV-2/SARS-CoV remodels the interaction pattern between NSP1 and neighboring proteins (uS3 and uS5) and rRNA (h18) lining the exit tunnel. This finding provides a clear picture of how SARS-CoV-2 invades human cells. However, the effect of NSP1 binding to 40S ribosome on mRNA translation has not been theoretically studied.

In this study, we used steered molecular dynamics (SMD) and alchemical simulations (Supporting Information (SI)), to explain how NSP1 binds to the 40S ribosome and inhibits the mRNA translation process. We found that the presence of NSP1 significantly increased the binding affinity of mRNA to 40S ribosome, which means that NSP1 binding to the mRNA channel inhibits its translation in the ribosome exit tunnel. Electrostatic mRNA-ribosome interactions have been found to play a key role in mRNA translation.

## Building two complexes

To study the effect of NSP1 on the binding affinity of mRNA to the ribosome, two complexes will be considered. One of these includes mRNA associated with the 40S ribosome in the absence of NSP1, and this complex will be referred to as mRNA-40S. The second complex, which will be referred to as mRNA-40S-NSP1, is similar to mRNA-40S, but in the presence of NSP1. Details on how mRNA-40S and mRNA-40S-NSP1 structures have been constructed are available from SI, but here we will mention the main steps. The structure of the 40S-NSP1 complex was retrieved from protein data bank (PDB) with PDB ID 6ZOJ.^30^ Then, the mRNA-40S-NSP1 structure was obtained by inserting the mRNA structure extracted from the 6HCJ PDB structure into 6ZOJ (Figure 1B).^31^ Finally, the mRNA-40S was obtained from mRNA-40S-NSP1 by removing NSP1. Our models include ribosomal proteins (rprotein) and ribosomal RNA (rRNA) of the 40S ribosome.

Because the mRNA has been mechanically inserted into the complexes, they should be allowed to relax before running the SMD simulation. Since the systems are large they may not be equilibrated using only all-atom simulations forcing us to combine coarse-grained and all-atom simulations (see SI). First, we performed energy minimization, followed by a short 5 ns simulation in NVT and NPT ensembles, and then a 1000 ns of conventional coarse-grained molecular dynamics (CGMD) simulation for mRNA-40S and mRNA-40S-NSP1 complexes using the MARTINI force field^32,33^ and coarse-grained water model.^34^ The last snapshot of the CGMD simulation was converted to the all-atom structure and its energy was minimized by using the steepest-descent algorithm, followed by a short simulation of 3 ns in NVT and NPT ensembles. A 200 ns production conventional molecular dynamics (CMD) simulation was performed using the AMBER99SB force field^35^ and the water model TIP3P.^36^

The root-mean-square displacement (RMSD) with respect to the original structure of the two complexes fluctuates during CGSM (Figure S1A) but remains below 0.35 nm, indicating that the system is equilibrated. This conclusion does not change after a 200 ns all-atom simulation because the structure remains nearly the same (Figure S1B). This conclusion does not change after all-atom simulation for 200 ns because the structure remains almost the same (Fig. S1B). Thus, the structure obtained in this production run can be used as the initial structure for SMD simulation.

## Hydrogen bonds and non-bonded contacts

To explore the network of hydrogen bonds (HBs) and non-bonded contacts (NBCs) between mRNA and the 40S ribosome, as well as the 40S-NSP1, we used the most populated structure obtained by clustering snapshots generated from 200 ns CMD simulations, as described in SI. HB and NBC were analyzed using Ligplot package^37^ (Figures S2A and S2B). HBs and NBCs between mRNA and 40S of the mRNA-40S are 27 and 7, while they are 29 and 23 for the mRNA-40S-NSP1. Thus, the mRNA- 40S-NSP1 has 2 HBs more than the mRNA-40S, and its NBCs are approximately three times larger. The difference in HBs and NBCs of the two complexes suggests that the binding affinity of mRNA to the 40S-NSP1 is higher than that of mRNA to the 40S ribosome. However, to confirm this conclusion, SMD simulations and more accurate alchemical calculations will be carried out. Moreover, HBs and NBCs between mRNA and NSP1 are 10 and 8 for the mRNA-40S-NSP1 complex, suggesting that NSP1 may significantly contribute to the binding affinity between mRNA and 40S- NSP1.

## Binding affinity of mRNA to 40S ribosome: SMD simulations

Details and setup of SMD simulations are described in SI (Figure S3). The 10 most representative structures obtained by clustering the snapshots collected in the 200 ns CMD run were used as initial conformations for the 10 SMD trajectories. Figure 2 shows the force, non-equilibrium work, and non-equilibrium binding free energy profiles of the mRNA-40S and mRNA-40S-NSP1 complexes. These results were averaged over 10 independent simulations.

**Figure 2:**
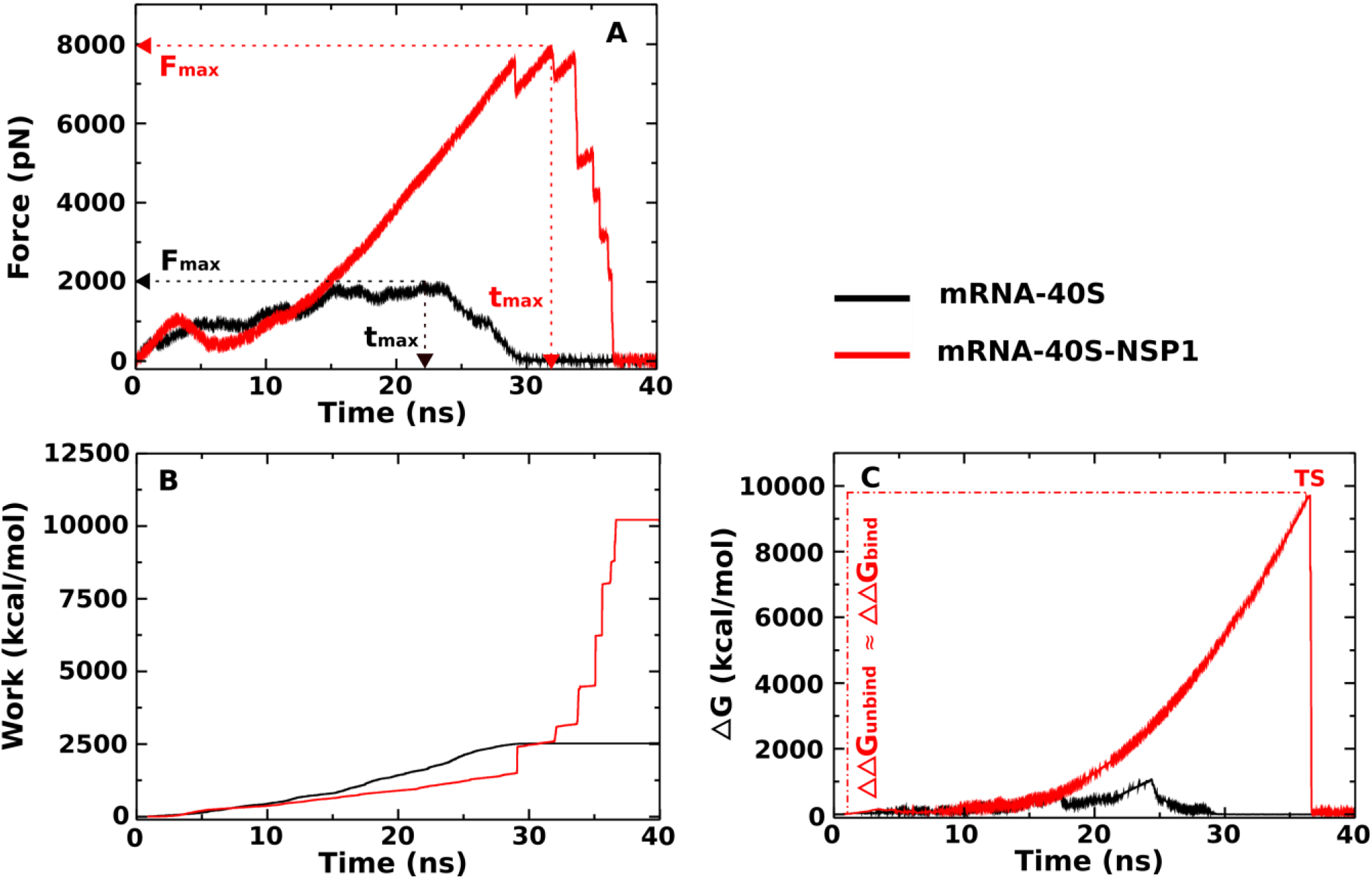
Time dependence of (A) force, (B) work, and (C) non-equilibrium energy profiles of the mRNA-40S and mRNA-40S-NSP1 complexes. The results were averaged over 10 independent SMD runs.

The force-time profile shows that mRNA binds to the 40S-NSP1 (*F_max_* = 7875.4±231.8 pN) are stronger than that to the 40S ribosome (*F_max_* = 1926.1±213.7 pN) (Figure 2A and Table 1). The time to reach the maximum force *t_max_* increases with increasing *F_max_*. The non-equilibrium work increased until the mRNA detached from the exit tunnel of the 40S ribosome and became saturated (Figure 2B). Defining the work performed by mRNA during the escape of the ribosome *W* as a saturated value at the end of the simulation, we obtained *W* = 2527.5±73.5 and 10175.7±147.7 kcal/mol for the mRNA- 40S and the mRNA-40S-NSP1, respectively (Table 1).

**Table 1:** Rupture force (*F_max_*), rupture time (*t_max_*), and non-equilibrium work (*W*), non-equilibrium binding energy barriers (ΔΔ*G*_bind_ and ΔΔ*G*_unbind_) were obtained from 10 independent SMD trajectories of the mRNA-40S and mRNA-40S-NSP1. The errors represent standard deviations.

|  | **mRNA-40S** | **mRNA-40S-NSP1** |
| --- | --- | --- |
| $F_{max}$ (pN) | 1926.1±213.7 | 7875.4±231.8 |
| $t_{max}$ (ps) | 22169.5±2013.1 | 31930.5±1767.6 |
| $W$ (kcal/mol) | 2527.5±73.5 | 10175.7±147.7 |
| $\Delta\Delta G_{unbind}$ (kcal/mol) | 1097.3±31.1 | 9737.3±87.1 |
| $\Delta\Delta G_{bind}$ (kcal/mol) | 1097.7±32.0 | 9738.7±87.7 |

The non-equilibrium binding free energy (Δ*G*) for the two complexes was also computed using equation S3 in SI. We define Δ*G*_bound_= Δ*G*(*t*_0_ =0) ≈ 0 kcal/mol as the free energy of the bound state, and the unbound state occurs at the end of the simulation with Δ*G*_unbound_ = Δ*G*(*t*_end_) ≈ 0 kcal/mol. The binding and unbinding free energy barriers are then calculated as ΔΔ*G*_bind_ = Δ*G*_TS_ – Δ*G*_unbound_ and ΔΔ*G*_unbind_ = Δ*G*_TS_ - Δ*G*_bound_, where Δ*G*_TS_ is the maximum free energy corresponding to the transition state (Figure 2C). ΔΔ*G*_bind_ and ΔΔ*G*_unbind_ are practically the same for each complex, ΔΔ*G*_unbind_ = 1097.3±31.1 and 9737.3±87.1 kcal/mol, and ΔΔ*G*_bind_ = 1097.7±32.0 and 9738.7±87.7 kcal/mol for the mRNA-40S and the mRNA-40S-NSP1, respectively (Table 1). Thus, the results obtained for *F*_max_, *W*, ΔΔ*G*_unbind_ and ΔΔ*G*_bind_ show that mRNA binds to the 40S-NSP1 much more strongly than to the 40S ribosome, indicating that NSP1 binding to the 40S ribosome stalls the mRNA translation process. Our results are in good agreement with the experiments.^22,30^

## NSP1 binding to 40S ribosome reduces the electrostatic and vdW interaction energies between mRNA and 40S ribosome

Averaging over 10 independent SMD runs, we obtained the van der Waals (*ΔE*_vdW_), electrostatic (*ΔE*_elec_), and total non-bonded (*ΔE_total_* = *ΔE*_elec_ + *ΔE*_vdW_) interaction energies as a function of simulation time shown in Figure 3. Obviously, *ΔE_vdW_* was negative in the bound state, then it reached 0 kcal/mol in the unbound state for both complexes (Figure 3A). In contrast, *ΔE_elec_* was positive in bound and unbound states (Figure 3B). *ΔE*_elec_ is much larger than *ΔE*_vdW_ for the mRNA-40S and mRNA-40S-NSP 1 complexes, resulting in *ΔE_total_* > 0 (Figure 3C). Since the bound state exists before the rupture occurs (*t* < *t*_max_), the energy of this state was obtained by averaging over the time window [0, *t*_max_], which gives *ΔE*_elec_ = 125493.1±263.2 and 77238.4±217.9 kcal/mol, *ΔE*_vdW_ = −178.8±2.6 and −130.5±2.7 kcal/mol, and *ΔE_total_* = 125314.3±265.8 and 77107.9±220.6 kcal/mol for the mRNA-40S and the mRNA-40S-NSP1, respectively (Table 2). Thus, for both complexes, the electrostatic interaction prevails over the vdW interaction. The positive value of *ΔE_total_* is due to repulsion between negatively charged of the 40S ribosome (−1215e), mRNA (−18e), and NSP1 (−3e) (Table S1). Nevertheless, this result also shows that NSP1 binding reduces the interaction between mRNA and 40S ribosome.

**Figure 3:**
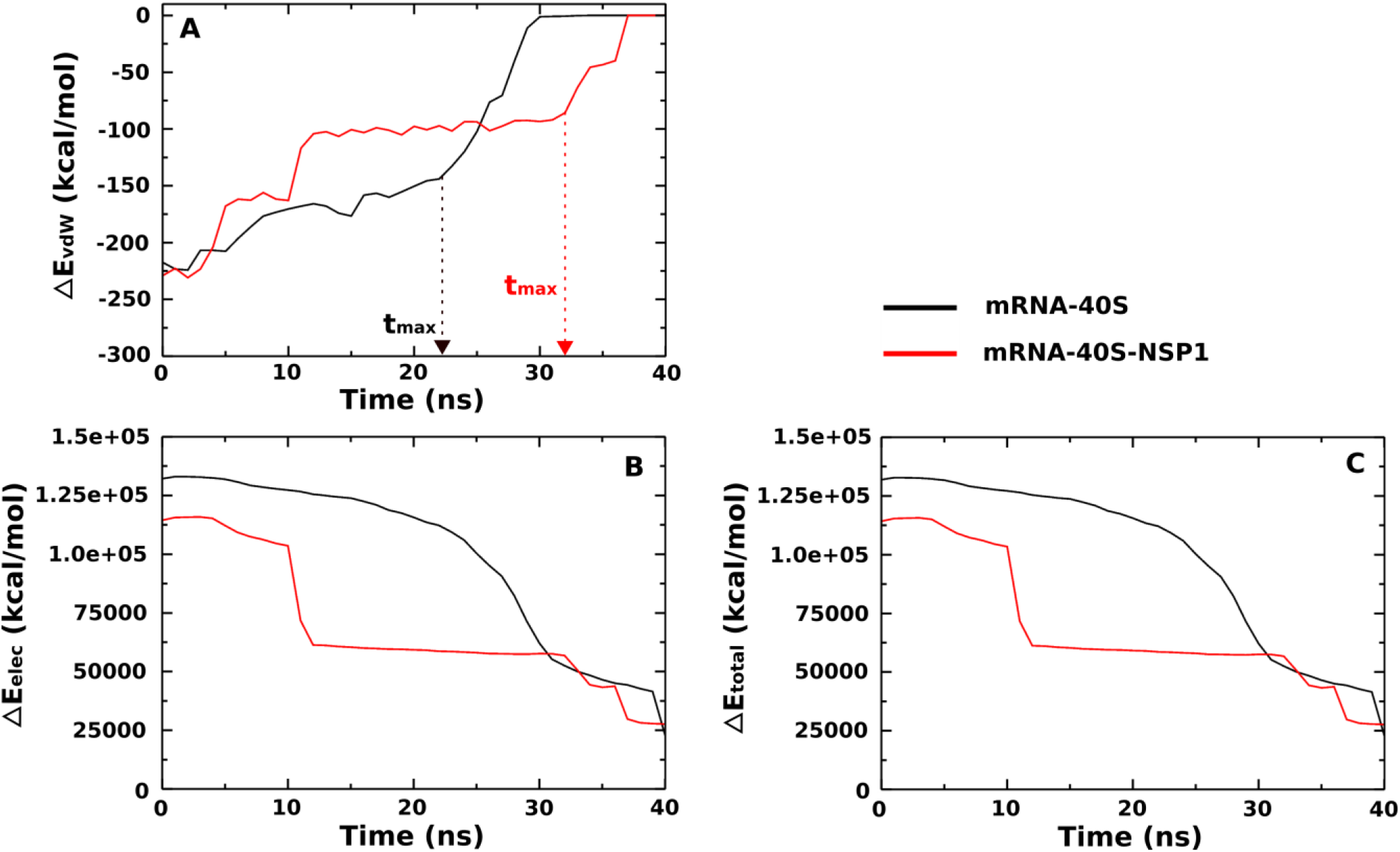
Time dependence of (A) vdW interaction energy, (B) electrostatic interaction energy, and (C) total non-bonded interaction energy of the mRNA-40S (black) and mRNA-40S-NSP1 (red). The results were averaged over 10 independent SMD runs.

**Table 2:** Non-bonded interaction energies (kcal/mol) of the mRNA-40S, and mRNA-40S-NSP1. The results were obtained for a [0-*t*_max_] time window and averaged over 10 SMD trajectories. The errors represent standard deviations.

|  | **mRNA-40S** | **mRNA-40S-NSP1** |
| --- | --- | --- |
| $\Delta E_{elec}$ | 125493.1±263.2 | 77238.4±217.9 |
| $\Delta E_{vdW}$ | -178.8±2.6 | -130.5±2.7 |
| $\Delta E_{total}$ | 125314.3±265.8 | 77107.9±220.6 |

## Water molecules stabilize the systems

Since the total interaction energy *ΔE_total_* obtained in the previous section is positive for both complexes, an important question emerges is whether these complexes are stable? To answer this question we will take into account water molecules. Again, *ΔE_total_* was calculated by averaging over 10 SMD trajectories in the time window [0, *t*_max_]. We obtained the total energy of −288365.7±224.1, and −305173.1±277.7 kcal/mol for the mRNA-40S and the mRNA-40S-NSP1, respectively (Table S2), which implies that these complexes are stabilized by water molecules.

## Important NSP1 residues

The per-nucleotide energy of mRNA and ribosome RNA (rRNA) and the per-residue energy of ribosome protein (rprotein) and NSP1 are shown in Figures 4A and 4B for both complexes. They were obtained by averaging over 10 SMD trajectories in the time window [0, t_max_], The energy of mRNA per nucleotide is much higher than that of rRNA, rprotein, and NSP1, the interaction of mRNA with all proteins and rRNA was taken into account, while for rRNA, rprotein, and NSP1, only the interaction with mRNA was considered. The total energy of nucleotides and amino acids in the binding region of mRNA-40S-NSP1 (77747.8 kcal/mol) is much more less than that of the mRNA-40S (110423.6 kcal/mol). This result is consistent with the result obtained for the entire system, including the binding region, according to which NSP1 reduces the interaction between mRNA and the 40S ribosome upon binding to the mRNA channel.

**Figure 4:**
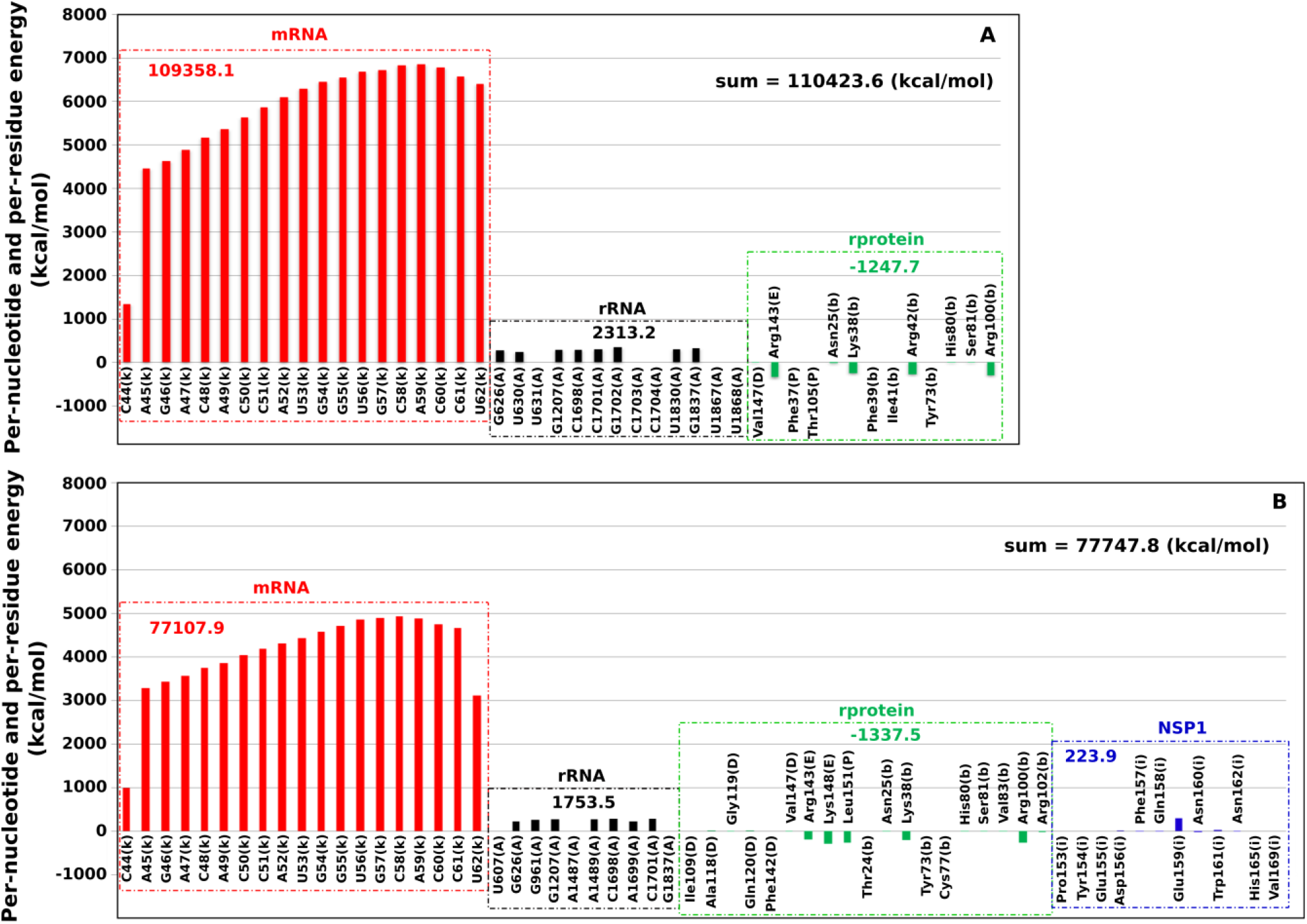
The interaction energy (electrostatic and vdW) per nucleotide and per residue at the binding regions of (A) mRNA-40S and (B) mRNA-40S-NSP1. The numbers next to the profiles refer to the total energy (sum over all interacting residues or nucleotides) measured in kcal/mol. For example, for mRNA in (A) the sum over all nucleotides and residues in the binding site is 110423.6 kcal/mol, while in (B) it is only 77747.8 kcal/mol. The results were averaged over 10 independent SMD runs.

Importantly, the contribution of each NSP1 residue at the binding region to the binding energy is Asp156 = 3.5, Phe157 = −17.3, Gln158 = −20.6, Glu159 = 281.9, Asn160 = −27.4, Trp161 = 13.6, and Asn162 = −9.8 kcal/mol (Figure 4B). Since the interaction energy of Phe157, Gln158, Asn160, and Asn162 is negative, these residues stabilize the system, while at a positive interaction energy, Asp156, Glu159, and Trp161 make the complex less stable. Despite the fact that the NSP1 total energy of interaction with mRNA is positive (223.9 kcal/mol), its presence makes the complex more stable by reducing the binding energy of mRNA, and the interaction energy of rRNA, and rprotein with mRNA. Taken together, mRNA translation at the 40S ribosome of the host immune system is controlled by electrostatic interactions and can be stalled by NSP1. The NSP1 residues Asp156, Phe157, Gln158, Glu159, Asn160, Trp161, and Asn162 play a key role as they are at the interface with mRNA.

## Binding free energy of mRNA to the 40S ribosome: Alchemical simulations

Since SMD at high pulling speeds only allows estimation of relative binding affinity, in order to evaluate the effect of NSP1 on the absolute binding affinity of mRNA to the 40S ribosome, alchemical free energy calculations were performed using the coarse grained MARTINI model. Note that there are many MD-based methods for estimating the binding free energy, but we have chosen alchemical modeling as this method is among the best (see SI). Alchemical free energy calculations are based on the thermodynamics cycle (Figure S4) allowing for calculating 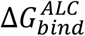 using Equation S5. For alchemical transformations, a set of *λ* values was used in the range from *λ* = 0 to *λ* = 1, where *λ* = 0 and *λ* = 1 correspond to a system with and without full interaction, respectively. To get the optimal set, 30, 30, and 20 windows of *λ* values were selected for mRNA-40S, mRNA-40S-NSP1, and mRNA, respectively (see SI). For each value of *λ*, a 1000 ns simulation was performed.

Figure S5 shows the time dependence of the root-mean-square deviation (RMSD) of mRNA, mRNA- 40S, and mRNA-40S-NSP1 at *λ* = 0, indicating that these systems have reached equilibrium after about 200 ns. Therefore, we calculated the binding free energy of mRNA to 40S ribosome and 40S-NSP1 for three time windows of [200-500ns], [200-800ns], and [200-1000ns] (Table 3). Within error bars, the results for the three windows are similar, implying that the data have been equilibrated. Hence, the result obtained in the [200-1000ns] window will be used. For mRNA-40S, the binding free energy 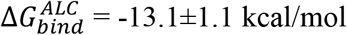, which is very close to the experimental value of −10.7±0.1 kcal/mol.^38^ For the mRNA-40S-NSP1, we obtained 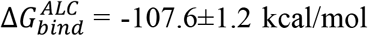, which is much smaller than that of the mRNA-40S. Overall, this finding is in line with SMD simulations, according to which the presence of NSP1 dramatically increases the binding affinity of mRNA to the ribosome, providing evidence that the translation process of mRNA can be completely prevented by NSP1, as observed in experiments.^22,30^

**Table 3:** The binding free energies (kcal/mol) of the mRNA-40S and mRNA-40S-NSP1 complexes were estimated from the alchemical molecular dynamics simulation using the MARTINI coarse-grained model.

|  |  | mRNA-40S | mRNA-40S-NSP1 |
| --- | --- | --- | --- |
| Experiment<sup>38</sup> |  | **-10.7±0.1** | NA |
| Our simulation ($\Delta G_{bind}^{ALC}$) | 200-500ns | -12.2±0.9 | -107.3±1.4 |
|  | 200-800ns | -12.7±1.2 | -108.2±1.2 |
|  | 200-1000ns | **-13.1±1.1** | **-107.6±1.2** |

## Conclusion

In conclusion, combining SMD and alchemical simulations, the association of mRNA with 40S ribosome in the absence and presence of NSP1 was studied. Our SMD results showed that mRNA binds to 40S-NSP1 much more strongly than to 40S ribosome, which is also in the line with the results obtained for the binding free energy using alchemical simulations and the MARTINI model. Our results are also in good agreement with the experimental data.^22,30^ The mRNA translation process was found to be driven by the electrostatic interaction between mRNA and 40S ribosome. After entering host cells, NSP1 may bind to the 40S ribosome and inhibit the translation process. Our analysis showed that the NSP1 residues Asp156, Phe157, Gln158, Glu159, Asn160, Trp161, and Asn162 at the interface with mRNA play a key role in triggering translational arrest of the host immune system.

## ASSOCIATED CONTENT

### Supporting Information

**Figure S1**: Root mean square deviation (RMSD) as a function of simulation time for mRNA-40S (back), and mRNA-40S-NSP1 (red) complexes. Results were obtained from CGMD (A) and all-atom CMD (B) simulations. **Figure S2**: Hydrogen bond and non-bonded contacts networks of (A) mRNA-40S and (B) mRNA-40S-NSP1. These networks were obtained using the most populated structure among snapshots generated from the 200 ns CMD simulation. The name of nucleotide/amino acid of mRNA (red), rRNA (black), rprotein (dark-green), and NSP1 (blue) is also displayed. **Figure S3**: Initial (left) and final (right) conformations from steered molecular dynamics simulation of the extraction of mRNA (red) from 40S ribosomal subunit (green-cyan: ribosomal protein (rprotein), and wheat: ribosomal RNA (rRNA)). **Figure S4**: An example of a thermodynamics cycle to calculate binding free energy between mRNA and 40S-NSP1 using alchemical simulation. State A (*λ* = 0) describes full interaction between mRNA and 40S-NSP1 while state B (*λ* = 1) presents mRNA (dummy) of no interaction to 40S-NSP1. The structures displayed in rRNA (wheat), rprotein (green-cyan), dummy (gray), mRNA (red), and NSP1 (blue). Alchemical free energy calculations are used in the MARTINI coarse-grained model. **Figure S5**: Root-mean-square deviation (RMSD) as a function of simulation time of mRNA (black), mRNA-40S (red), and mRNA-40S-NSP1 (green) at *λ* = 0 in the alchemical free energy calculations using the MARTINI coarse-grained model. The arrow indicates the time (200 ns) when the system reaches equilibrium. **Table S1**: Total charge of 40S ribosome, NSP1 and mRNA. **Table S2**: Total non-bonded energy of the mRNA-40S and mRNA- 40S-NSP1 complexes with water and ions (kcal/mol). The results were averaged over 10 independent SMD runs for the time window [0, *t*_max_].

### Notes

The authors declare no competing financial interest.

## Acknowledgments

This research was supported by Narodowe Centrum Nauki in Poland (Grant 2019/35/B/ST4/02086) and in part by PLGrid Infrastructure and the supercomputer centre TASK in Gdansk, Poland.

