## Supplementary material for "Binding of SARS-CoV-2 non-structured protein 1 to 40S ribosome inhibits mRNA translation": v1_SI_Li: v1_SI_Li.docx

**Table of Content**

Supporting Methods..................................................................................................S2

System Preparation.............................................................................................S2

Estimation of Binding Affinity Using Molecular Dynamics Simulations.........S2

Conventional and Steered Molecular Dynamics Simulations.....................S2

Alchemical Molecular Dynamics Simulations............................................S4

Supporting Figures....................................................................................................S6

Figure S1............................................................................................................S6

Figure S2............................................................................................................S7

Figure S3.............................................................................................................S8

Figure S4.............................................................................................................S9

Figure S5............................................................................................................S10

Supporting Tables.....................................................................................................S11

Table S1.............................................................................................................S11

Table S2.............................................................................................................S12

Supporting References..............................................................................................S13

**Supporting Methods**

*System preparation*

The three-dimensional structure of NSP1 associated with the 40S ribosome, which will be referred to as the 40S-NSP1, was taken from the Protein Data Bank (PDB) with ID 6ZOJ.^1^ Then, the complex of mRNA bound to 40S-NSP1 was constructed from the superposition of two PDB structures 6ZOJ and 6HCJ,^1,2^ where the mRNA structure was extracted from 6HCJ.^2^ Based on the cryo-EM structure of NSP1 in a complex with the 40S ribosome, we built a model system that includes ribosomal proteins (rprotein) of the 40S ribosome resolved in a cryo-EM structure, NSP1, ribosomal RNA (rRNA), 165 Mg^2+^, and 2 Zn^2+^ions, hereinafter termed SARS-CoV-2 in explicit water solvation. The mRNA-40S-NSP1 complex is displayed by using the PYMOL package^3^ (Figure 1B).

*Estimation of binding affinity using molecular dynamics simulations*

Several methods have been proposed to evaluate the binding free energy of protein-ligand, protein-protein, protein-DNA/RNA complexes,^4^ such as thermodynamics integration (TI),^5^ free energy perturbation (FEP),^6^ molecular mechanics with Poisson-Boltzmann or generalized Born and surface area (MM-PBSA and MM-GBSA),^7^ linear interaction energy (LIE),^8^ steered molecular dynamics (SMD).^9-11^

The alchemical method referred to as TI or FEP, which is based on MD simulation in explicit solvent, is known to be the most accurate method due to its high level of theoretical rigor, and is also accessible within existing computational powers for relatively large systems. This method is based on the non-physical thermodynamics cycle, where the binding free energy is calculated as the sum of multiple steps during which a ligand/protein/DNA/RNA is “inserted” or “removed” from different states, including bound and unbound states.^12-14^ The alchemical free energy calculation has been successful in determining free energy difference in many situations such as the partition of a compound between different media, the binding affinity of protein-ligand, protein-protein, protein-DNA/RNA complexes with mutations at their interface.^15^ The free energy difference can be calculated using alchemical routes of relative or absolute binding free energies.^16^

In this study, we conducted all-atom SMD simulations and a set of alchemical free energy calculations using the MARTINI coarse-grained model to evaluate the binding affinity of mRNA to the 40S ribosome and the 40S-NSP1.

*Conventional and steered molecular dynamics simulations*

Before starting SMD and alchemical simulation, we performed a 1000 ns coarse-grained molecular dynamics (CGMD) simulation for both the mRNA-40S and mRNA-40S-NSP1 complexes. The final frame of the CGMD simulations was then converted to an all-atom structure using the backward script.^17^ The resulting all-atom structure was used to run all-atom conventional molecular dynamics (CMD) simulations for 200 ns for mRNA-40S and mRNA-40S-NSP1 complexes to generate initial configurations for SMD and alchemical simulations.

For CGMD simulations, the complexes were placed in a dodecahedron box with a distance between the solute and the box of 1.2 nm. Since mRNA taken from the 6HCJ PDB structure was inserted into the 6ZOJ PDB structure, the energy of the system was first minimized, and then followed by a 5 ns NVT and NPT simulations. A 1000 ns CGMD simulation was then performed for these systems using MARTINI force field^18-20^. The root mean square deviation (RMSD) versus time from the CGMD simulation is shown in Figure S1A. Obviously, the two complexes have reached equilibrium, but it remains unclear whether the CG structure is stable in all-atom models. To test this the last snapshot was converted to the all-atom structure, which was used as the initial structure for the all-atom CMD simulation. In this simulation, the complexes were placed in a dodecahedron box with a distance of 1.2 nm between the solute and the box. The system energy was then minimized using the steepest descent algorithm followed by a short 3 ns CMD simulation in the NVT and NPT ensembles using the AMBER99SB force field and the TIP3P water model. Finally, a 200 ns production CMD simulation was conducted using the leap-frog algorithm.^21^

As can be seen from the RMSD time dependence (Fig. S1B), the complexes are also stable in all-atom models. Applying the clustering analysis to the collected snapshots from a 200 ns all-atom CMD run, we obtained 10 representative structures that will be used as the initial structure to run 10 independent SMD simulations. These structures were also used for alchemical simulations.

SMD simulations^9-11^ were then carried out to pull mRNA from the ribosomal 40S exit tunnel for cases with and without NSP1. Pulling speeds *v* = 0.5 nm/ns was used resulting in a simulation time of 40 ns. This value of *v* is about ten orders of magnitude larger than in the experiment, but as shown in previous works,^22-24^ this choice does not influence relative binding affinities, *i.e.*, it can be used to discern strong binders from weak ones. A rectangular box with the dimension of 32×25×55 nm^3^ was used for all systems. The complexes were immersed in 0.15 M sodium chloride solution and counter ions were added to neutralize the system. To pull the mRNA out of the exit tunnel, an external force is applied to the dummy atom connected to the 5'-mRNA (O5’ atom), through a spring with a stiffness of k. The complexes were rotated so that the exit direction was parallel to the z-axis (Figure S3).

The force experienced by a stretched molecule is calculated as follows:

$F=k\left( \Delta z-vt \right)$	(S1)

where *k* is the stiffness of the spring, *v* is the pulling speed, *∆z* is the displacement of the atom connected to the spring in the direction of pulling. The spring constant *k* was set to 600 kJ/(mol.nm^2^) (≈ 1020 pN/nm), which is a typical value used in atomic force microscopy (AFM) experiments.^25^

Using the force-displacement profile obtained from SMD simulations, the non-equilibrium work (*W*) can be calculated using the trapezoidal rule:

$W=\int Fdz=\sum_{i=1}^{N} \frac{F_{i+1}+F_{i}}{2}\left( z_{i+1}-z_{i} \right)$	(S2)

where *N* is the number of simulation steps, *F_i_* and z*_i_* are the force defined by equation S1 and the position at step *i*, respectively.

To estimate the binding free energy (*∆G*), we used Jarzynski’s equality^26,27^ extended to the case when the applied external force grows at a constant speed *v*:

$exp\left( \frac{-\Delta G}{k_{B}T} \right)=\left\langle exp\left( \frac{W_{t}-\frac{1}{2}k\left( z_{t}-vt \right)^{2}}{k_{B}T} \right) \right\rangle_{N}$	(S3)

Here is the average over *N* trajectories, *z_t_* is the time dependent displacement, and *W_t_* is the non-equilibrium work at time *t* defined by equation S2.

From equation S3, the equilibrium free energy can be extracted if the number of simulations is large enough and the pulling sufficiently slow. However, since the pulling is not slow enough in our case and the number of SMD runs is limited, we can only estimate the non-equilibrium binding and unbinding barriers separating the transition state (TS) from the bound state at *t*=0 and the unbound state at *t*_end_.^28^

Both CMD and SMD simulations were performed using the AMBER99SB force field^29^ implemented in the GROMACS 2016 package.^30^ The v-rescale^31^ and Parrinello-Rahman^32^ algorithms were used to maintain the temperature at 310 K and the isotropic pressure at 1 bar, respectively. The water model TIP3P^33^ was used for all systems. Bond lengths were constrained by the linear constraint solver (LINCS) algorithm^34^ allowing a time step of 2 fs. The electrostatic and van der Waals interactions were calculated using a cutoff of 1.4 nm, and the non-bonded interaction pair-list was updated every 10 fs. The Particle Mesh Ewald algorithm^35^ was used to treat long-range electrostatic interactions. Periodic boundary conditions were applied in all directions.

*Alchemical molecular dynamics simulations*

The most populated structure from the 200 ns all-atom CMD simulations was chosen as the initial structure for the alchemical free energy calculation of mRNA-40S and mRNA-40S-NSP1. The standard coarse-grained MARTINI 2.2 force field, which was developed for modeling biological systems such as biological membranes, proteins, nucleotides, *etc.*^18-20^, was used to calculate the binding free energy by this approach. This force field is accurate enough to describe the ligand-protein, protein-protein, protein-DNA/RNA, and protein-liquid interaction in an aqueous medium.^18-20,36,37^ The MARTINI water model^38^ was used with a minimum distance between water beads of 1.0 nm. The system was neutralized by adding sodium chloride salt solution. The temperature was set to T = 300 K using a v-rescale thermostat,^31^ and pressure was set to p = 1.0 bar with a Parrinello-Rahmanbarostat.^32^ The LINCS algorithm^34^ was used to constrain the length of all bonds.

To evaluate the free energy of mRNA binding to the 40S ribosome with and without NSP1, we created the thermodynamic cycle described in Figure S4. From the thermodynamic cycle, we have:

${\Delta G}_{bind}^{ALC}-\Delta G={\Delta G}_{complexation}-{\Delta G}_{solvation}$	(S4)

∆G≡0 as it is related to non-interacting (λ=1) mRNA being dummy and dummy-40S-NSP1. Then the binding free energy has the following form:

${\Delta G}_{bind}^{ALC}={\Delta G}_{complexation}-{\Delta G}_{solvation}$	(S5)

For alchemical transformations, we used a set of λ values ranging from λ = 0 to λ = 1, where λ = 0 and λ = 1 correspond to a system with and without full interaction, respectively. To obtain the optimal set, 30, 30, and 20 windows of λ values were selected for the mRNA-40S, mRNA-40S-NSP1, and mRNA, respectively. Thus, a total of 80 windows were used for alchemical calculations of free energy.

Optimal sets of λ-values were generated using the available script at <https://gitlab.com/KomBioMol/converge_lambdas.>^39^ For each window, simulations were run for 1000ns to ensure that the complexes reached equilibrium. Free energy changes were estimated using the Bennett acceptance ratio (BAR).^40^ The binding free energy was then calculated from the thermodynamics cycle (Figure S4).

**Supporting Figures**

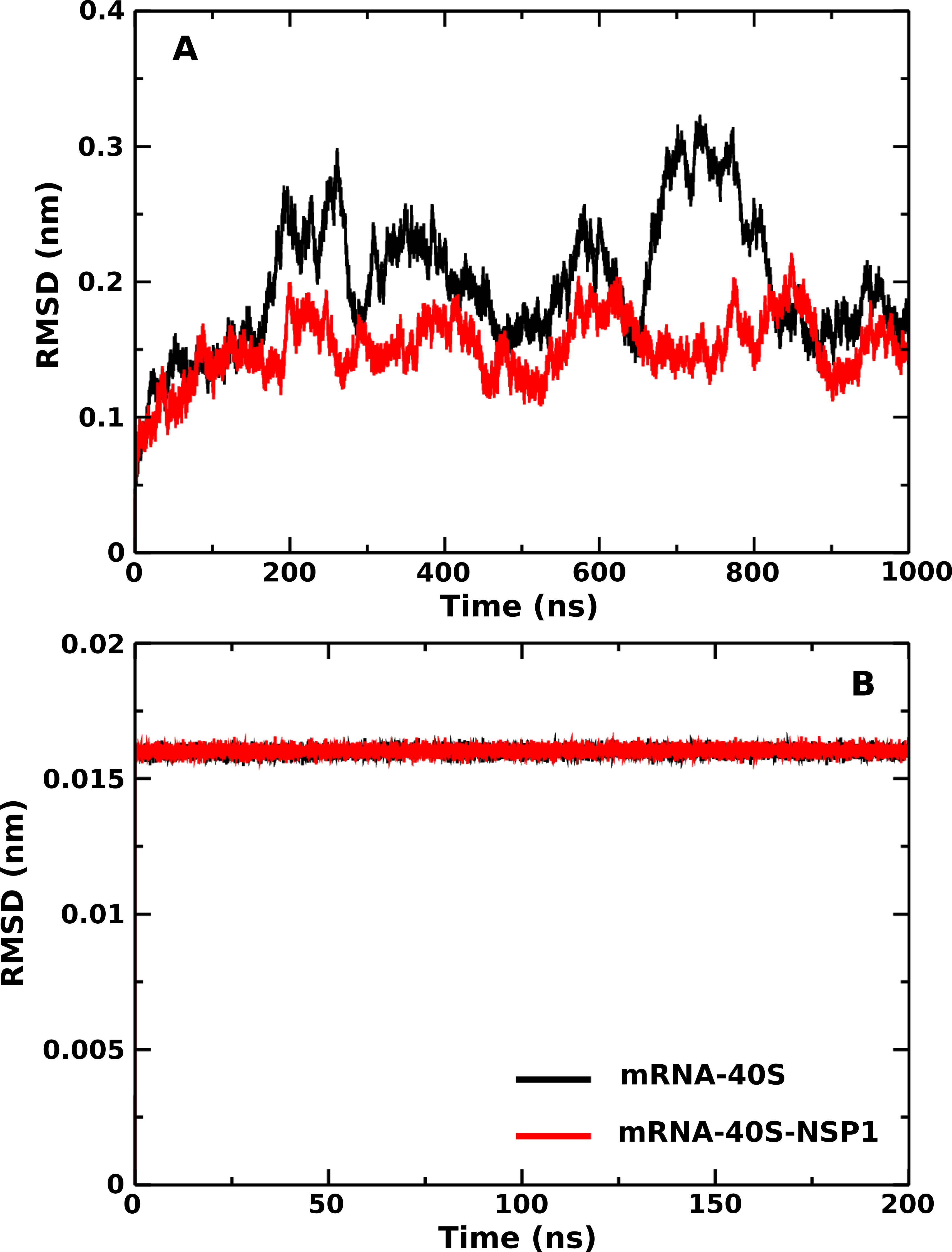

**Figure S1**: Root mean square deviation (RMSD) as a function of simulation time for mRNA-40S (back), and mRNA-40S-NSP1 (red) complexes. Results were obtained from CGMD (A) and all-atom CMD (B) simulations.

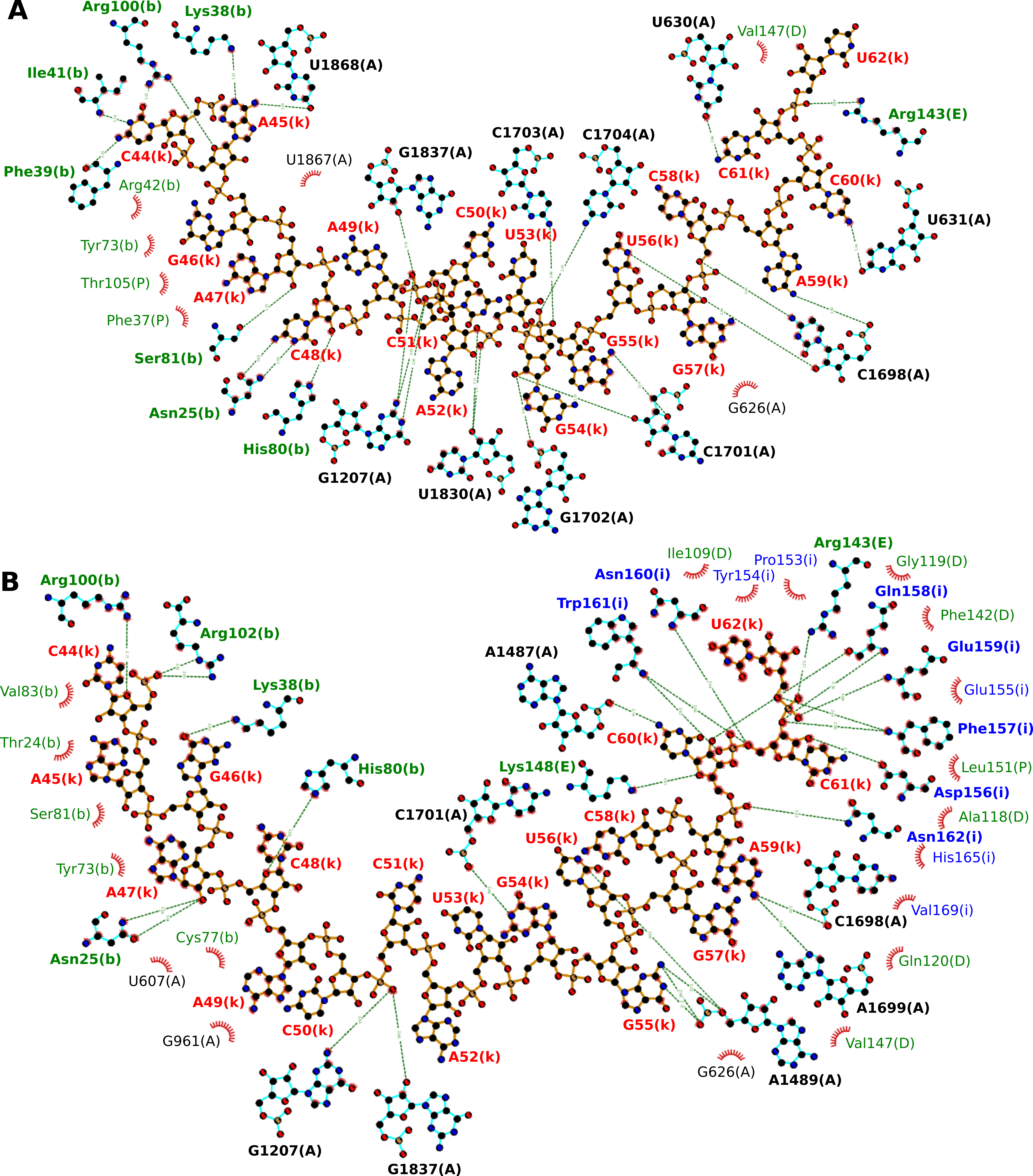

**Figure S2**: Hydrogen bond and non-bonded contacts networks of (A) mRNA-40S and (B) mRNA-40S-NSP1. These networks were obtained using the most populated structure among snapshots generated from the 200 ns CMD simulation. The name of nucleotide/amino acid of mRNA (red), rRNA (black), rprotein (dark-green), and NSP1 (blue) is also displayed.

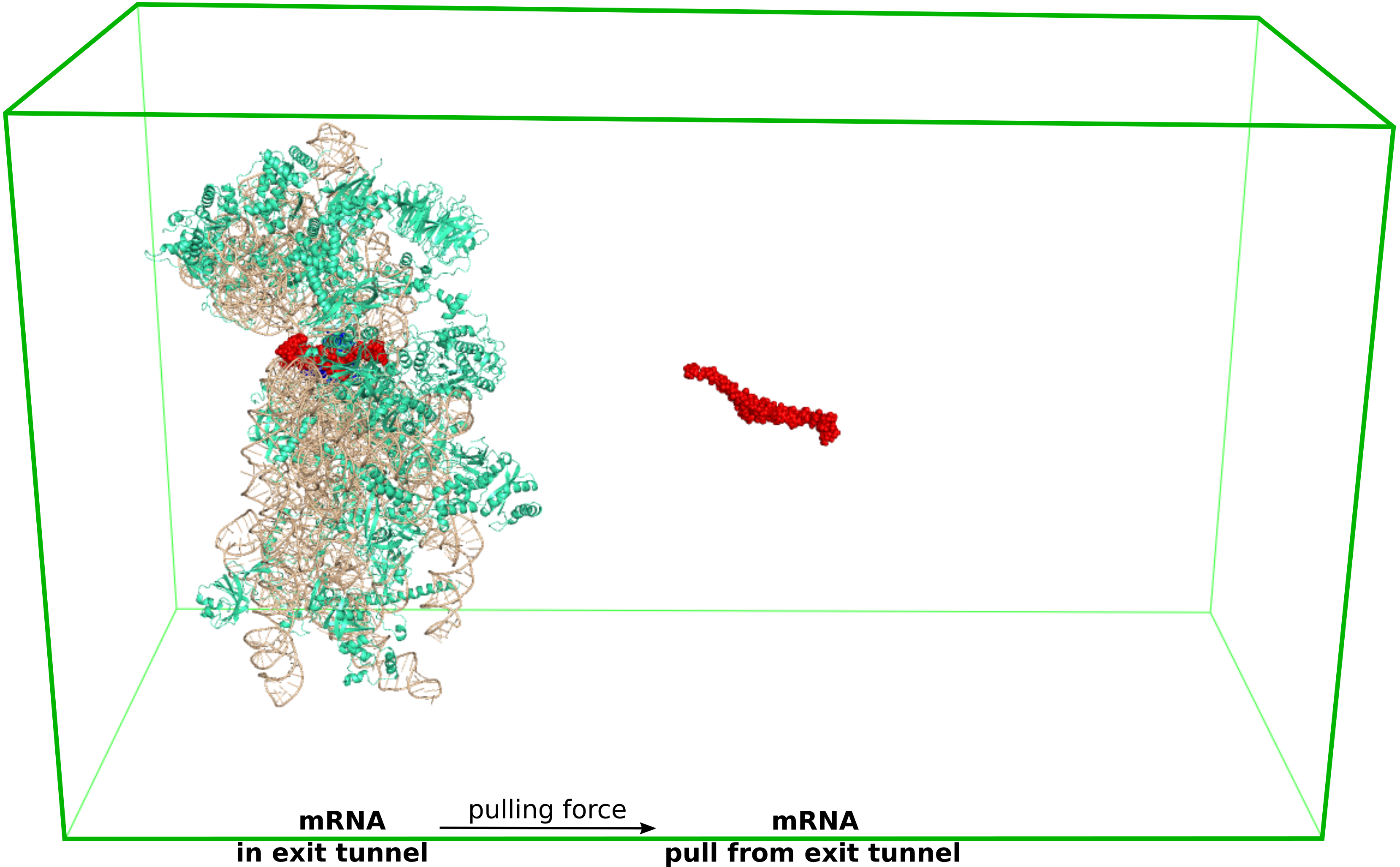

**Figure S3**: Initial (left) and final (right) conformations from steered molecular dynamics simulation of the extraction of mRNA (red) from 40S ribosomal subunit (green-cyan: ribosomal protein (rprotein), and wheat: ribosomal RNA (rRNA)) and NSP1.

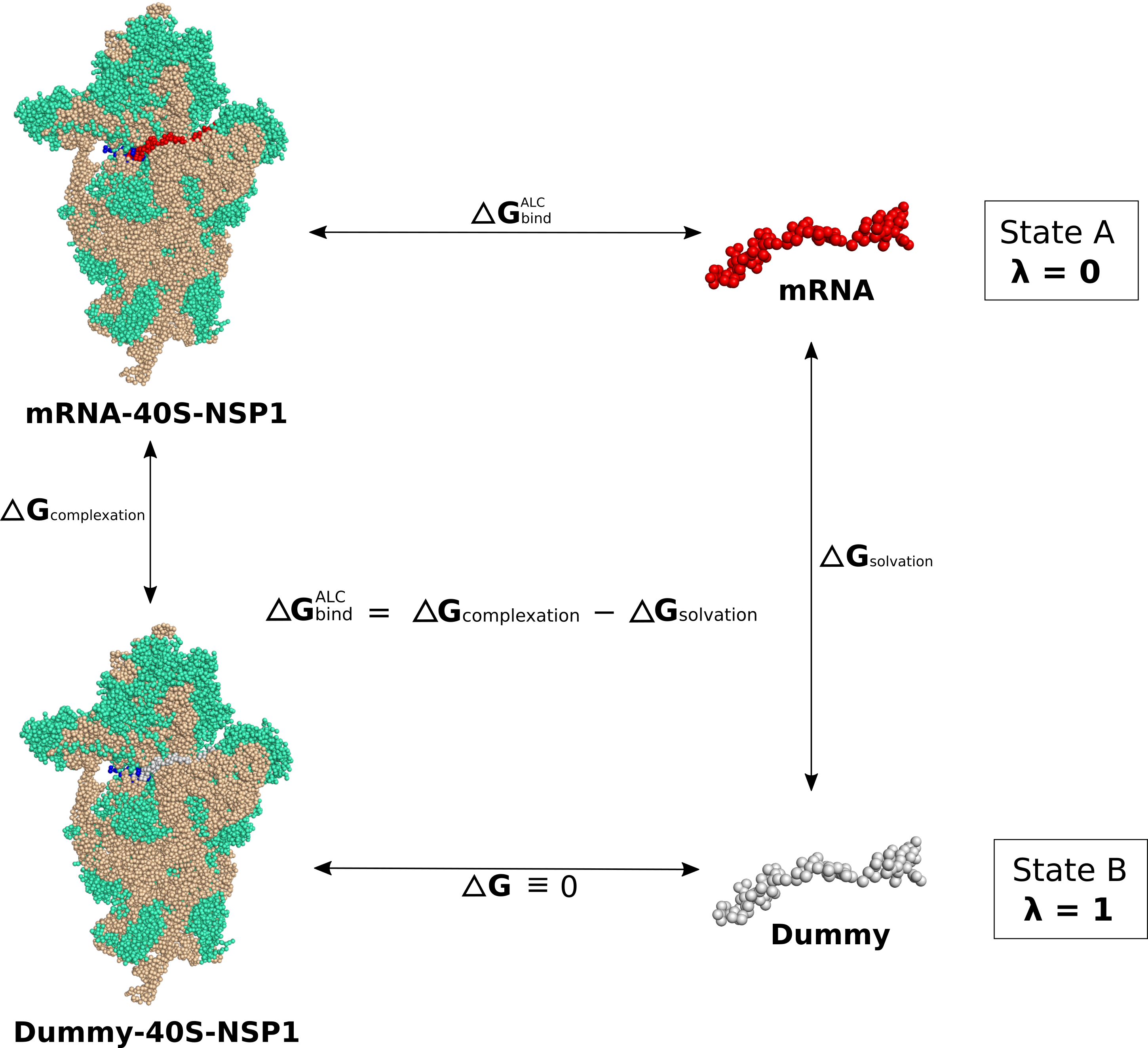

**Figure S4**: An example of a thermodynamics cycle to calculate binding free energy between mRNA and 40S-NSP1 using alchemical simulation. State A (λ = 0) describes full interaction between mRNA and 40S-NSP1 while state B (λ = 1) presents mRNA (dummy) of no interaction to 40S-NSP1. The structures displayed in rRNA (wheat), rprotein (green-cyan), dummy (gray), mRNA (red), and NSP1 (blue). Alchemical free energy calculations are used in the MARTINI coarse-grained model.

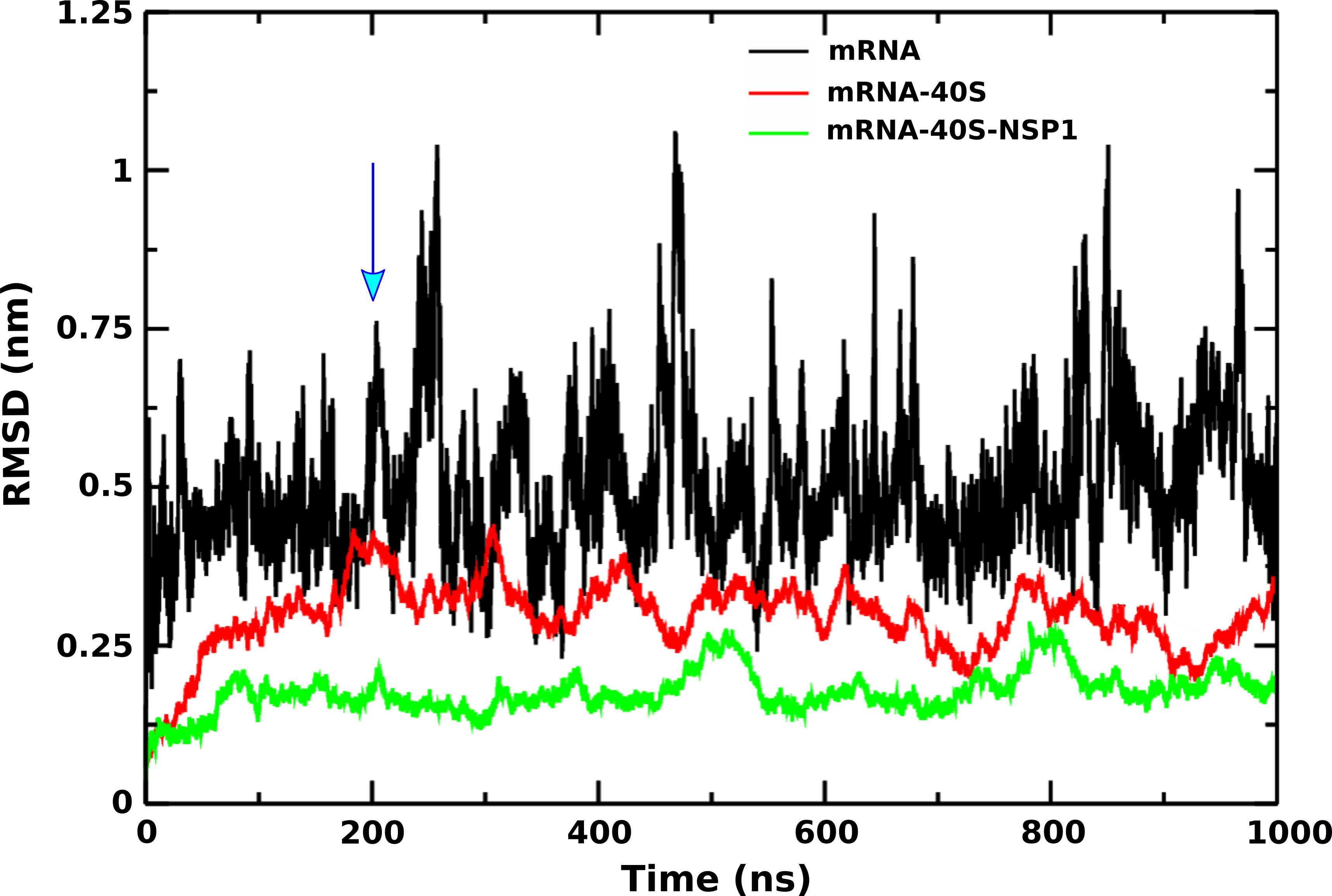

**Figure S5**: Root-mean-square deviation (RMSD) as a function of simulation time of mRNA (black), mRNA-40S (red), and mRNA-40S-NSP1 (green) at λ = 0 in the alchemical free energy calculations using the MARTINI coarse-grained model. The arrow indicates the time (200 ns) when the system reaches equilibrium.

**Supporting Tables**

**Table S1**: Total charge of 40S ribosome, NSP1 and mRNA.

| **Structure** | **Total charge (e)** |
| --- | --- |
| 40S ribosome | -1215 |
| NSP1 | -3 |
| mRNA | -18 |

**Table S2**: Total non-bonded energy of the mRNA-40S and mRNA-40S-NSP1 complexes with water and ions (kcal/mol). The results were averaged over 10 independent SMD runs for the time window [0, *t*_max_].

| **Complex** | **Total non-bonded energy** |
| --- | --- |
| mRNA-40S with water and ions | -288365.7±224.1 |
| mRNA-40S-NSP1 with water and ions | -305173.1±277.7 |
